# Consistent gut bacterial microbiota in European sea bass fed aquafeeds containing sustainable plant and invasive fish-based ingredients

**DOI:** 10.64898/2026.06.26.733563

**Authors:** Eleni Nikouli, Antigoni Vasilaki, Ioannis Nengas, Anna Tampou, Eleni Mente, Konstantinos Kormas

**Author notes:** Equal contribution (Eleni Nikouli, Antigoni Vasilaki). Department of Fisheries and Aquaculture, School of Agricultural Sciences, University of Patras, Mesolonghi, Greece (Anna Tampou).

## Abstract

The aim of this study was to evaluate the impact of two sustainable dietary protein sources on the structure and composition of the gut microbiota in European sea bass (*Dicentrarchus labrax*) juveniles. These protein sources were incorporated to the aquafeeds containing (a) *Lupinus albus* meal, treated with either exogenous enzymes (Solid state hydrolysis-SSH) or fermented with *Saccharomyces cerevisiae* (Solid state fermentation, SSF) and (b) *Lagocephalus sceleratus* meal. In the first case (a), the control aquafeed simulated a standard commercial diet, containing soybean meal whereas in the rest of the diets soybean meal was partially or totally replaced by hydrolysed or fermented Lupin meal. In the second case (b) the fish were fed *Lagocephalus sceleratus* unprocessed fishmeal as well as treated at different temperatures to deactivate tetrodotoxin (TTX). A control diet with 30% commercial fish meal was also fed as a reference diet. Both diets in all inclusion levels did not cause any significant gut microbiota change, suggesting their neutral role in this aspect. However, the gut bacterial communities of the fish fed with 12.5% lupin meal inclusion, had increased amino acid biosynthetic pathways suggesting a beneficial effect.

## INTRODUCTION

The growing demand for aquaculture products has intensified the need for more sustainable aquafeed solutions, driving efforts to reduce reliance on conventional marine ingredients such as fishmeal and fish oil. This transition is motivated by concerns regarding the long-term availability of marine resources, environmental sustainability and the need to ensure global food security while maintaining marine biodiversity and without compromising the farmed fish performance, health and welfare. Consequently, considerable research has focused on the identification and evaluation of alternative feed ingredients—including plant-based proteins, microbial and algal biomass, insect meals—that can meet the nutritional requirements of farmed species while reducing the environmental footprint of aquaculture (Panteli et al., 2025; Serra et al., 2024). This merging of sustainability goals with feed innovation reflects an urgent and strategic response to the global challenges facing modern aquaculture.

Lupin has been identified as a promising alternative protein source due to its nutritional value, digestibility and positive effects on the performance of several aquaculture species (Malarvizhi et al., 2024; Szczepański et al., 2022). Lupin derived ingredients have been successfully incorporated in aquafeeds replacing up to 40-50% of fishmeal in a wide range of species including rainbow trout, Atlantic salmon, common carp, perciform fish (sparids, barramundi, tilapia), and shrimps (Szczepański et al., 2022). More recently, biotechnologically processed lupin (*Lupinus albus*), either fermented with *Saccharomyces cerevisiae* or hydrolysed using exogenous enzymes, was evaluated as a successful replacement for soybean meal in diets for the European sea bass (*Dicentrarchus labrax*). Inclusion levels of up to 12.5% resulted in improved feed utilization, enhanced apparent digestibility coefficients of protein and lipid, and moderate immunostimulatory effects (Vasilaki et al., 2025a, 2025b). These findings highlight the potential of biotechnologically processed lupin as a sustainable and functional aquafeed ingredient for marine carnivorous fish. Despite the growing interest in lupin as feed ingredient, its effects on the gut microbiota of farmed fish remain relatively underexplored. To date, the relationship between dietary lupin inclusion and gut microbiota has been investigated in gilthead sea bream (*Sparus aurata*) (Silva et al., 2011) and in combination with other replacement ingredients in barramundi (*Lates calcarifer*) (Gupta et al., 2020) and European sea bass (Gatesoupe et al., 2016, 2014). Consequently, further research is needed to better understand the microbiome mediated effects of lupin-based diets in aquaculture species.

The quest for more sustainable aquafeeds does not stop in plant-based materials as the industrial scale of plant-based protein processing has high demands for natural resources, energy, chemicals, and highly specialized equipment (Aimutis and Shirwaiker, 2024). Although plant-derived ingredients have become essential components of modern aquafeeds due to sustainability, economic, and supply limitations associated with fishmeal and fish oil, their use in diets for marine carnivorous fish remains associated with several nutritional and functional challenges. The increasing dependence of the aquafeed industry on terrestrial agroecosystems has further raised concerns regarding long-term sustainability, competition with human food systems, and environmental footprint, thereby driving interest toward more resilient and consistent alternatives. Within a circular economy perspective, increasing attention has been given to the use of animal-based biomass and ingredients, particularly those of marine origin for marine carnivorous fishes. Animal by-products were initially considered among the most promising alternatives, however, in recent years, non-indigenous and invasive species are gaining momentum as potential feed resources (Eroldoğan et al., 2023). The interest in utilizing non-indigenous animal biomass has become so intense, that the term “invasivevorism”, referring to the human consumption of invasive species, has recently emerged and has even been positively evaluated in some contexts (Oficialdegui et al., 2026), despite ongoing concerns regarding safety, regulation and consumer acceptance. In aquafeeds, the exploitation of invasive species offers a potential “double dividend” by simultaneously contributing to the mitigation of ecological impacts while generating valuable protein source raw materials. Consequently, the use of invasive alien fish species has been proposed as a viable short-term approach for sustainable aquafeed production. In this context, the utilization of invasive, unwanted or previously unexploited fish species represents an attractive approach, as it may reduce reliance on conventional feed resources while simultaneously enhancing circularity in aquafeed raw material production (Ragaza et al., 2021).

The existing studies investigating invasive fish species as aquafeed ingredients for farmed fish remain relatively scarce. Positive growth and feed utilization responses have been reported in thinlip mullet (*Chelon ramada*) and Nile tilapia (*Oreochromis niloticus*) fed diets containing invasive blue crab (*Portunus segnis*) meal (Hachana et al., 2025). Similarly, Amazon sailfin catfish (*Pterygoplichthys pardalis*) meal has been successfully evaluated as a dietary ingredient for Nile tilapia (Darwis et al., 2026, 2025), while the invasive suckermouth catfish (*Hypostomus plecostomus*) has also been identified as a promising aquafeed ingredient (Islam et al., 2026). However, while these studies have primarily focused on growth performance and nutritional evaluation, the effects of invasive species-derived ingredients on the gut microbial communities of farmed fish remain largely unexplored (Darwis et al., 2026). Today, *Lagocephalus sceleratus*, despite being one of the most impactful invasive fish species in Greek waters (Vagenas et al., 2024) and throughout the Mediterranean Sea, as was previously predicted by (Coro et al., 2018), has not yet been evaluated with respect to its effects on gut microbiota/microbiome of farmed fish. Owing to its abundance, high biomass availability and protein rich content, *L. sceleratus* has attracted interest as a potential resource for feed application in the recent research project “LagoMeal” (https://lagomeal.gr; funded by the European Maritime, Fisheries and Aquaculture Fund), aiming at transforming a destructive, toxic invasive species into sustainable, high-protein aquafeed ingredient, following the inactivation of its endogenous tetrodotoxin. The current research gap regarding the response of the fish gut microbiota to diets containing biomass derived from invasive species is particularly important considering that the fish gut microbiome is now recognised as a key determinant of host nutrition, metabolism, immune function and overall health in both wild and farmed fish (Piazzon et al., 2025). Furthermore, the gut microbiome has emerged as a major target for dietary interventions aimed at improving fish performance, welfare and disease resistance (Gupta et al., 2024; Luna et al., 2022; Zhang et al., 2025).

In this study, we hypothesised that the restructuring of the gut microbiota following the dietary inclusion of fermented lupin or hydrolysed lupin or *Lagocephalus sceleratus* fishmeal in aquafeeds of the European sea bass (*Dicentrarchus labrax*) juveniles would result either in only minor alterations compared with a conventional aquafeed or, ideally, in shifts toward to a more beneficial for the host gut microbial community. The lupin-containing aquafeeds were improved by solid state fermentation (SSF) with *Saccharomyces cerevisiae* and by solid state hydrolysis (SSH) with a complex of exogenous enzymes. To our knowledge, this is the first study evaluating the effects of dietary inclusion of *Lagocephalus sceleratus* biomass on the gut microbiota of a farmed fish species.

## MATERIALS AND METHODS

### Fish feeding trials

This study consisted of three series of feeding trials evaluating different sustainable alternatives to both soybean meal and fish meal. The first two feeding trials investigated the dietary inclusion of *L. albus* meal (variety Tennis), processed using biotechnological approaches to enhance nutrient availability and digestibility. Specifically, lupin meal was subjected either to SSH using exogenous enzymes (Synergen, Alltech Inc, Nicholasville KY, USA) (Table 1) or to solid state fermentation (FRL) with *Saccharomyces cerevisiae* (Table 2). Different inclusion levels of the processed lupin meals were subsequently evaluated in experimental diets.

**Table 1.** Formulation (% of wet weight) and composition of the *L. albus* meal diets treated with exogenous enzymes.

|  | **Control** | **Lupin 1** | **Lupin 2** | **Lupin 3** |
| --- | --- | --- | --- | --- |
| **Fishmeal<sup>a</sup>** | 20.00 | 20.00 | 20.00 | 20.00 |
| **Krill meal<sup>b</sup>** | 3.00 | 3.00 | 3.00 | 3.00 |
| **Blood meal<sup>b</sup>** | 5.00 | 5.00 | 5.00 | 5.00 |
| **Wheat meal<sup>b</sup>** | 9.40 | 11.20 | 11.70 | 11.70 |
| **Wheat gluten<sup>b</sup>** | 10.00 | 11.00 | 11.00 | 11.00 |
| **Corn gluten<sup>b</sup>** | 15.00 | 15.00 | 15.00 | 15.00 |
| **Soy protein concentrate<sup>b</sup>** | 5.00 | 5.00 | 5.00 | 5.00 |
| **Soybean meal (44% CP)<sup>b</sup>** | **15.00** | **5.00** | **2.50** | **0.00** |
| **Hydrolysed Lupin meal<sup>c</sup>** | **0.00** | **7.50** | **10.00** | **12.50** |
| **Fish oil<sup>d</sup>** | 13.00 | 12.50 | 12.00 | 12.00 |
| **Monocalcium Phosphate<sup>b</sup>** | 2.00 | 2.00 | 2.00 | 2.00 |
| **Mineral premix<sup>e</sup>** | 0.20 | 0.20 | 0.20 | 0.20 |
| **Vitamin premix<sup>f</sup>** | 0.20 | 0.20 | 0.20 | 0.20 |
| **L-Lysine HCL<sup>b</sup>** | 1.00 | 1.20 | 1.20 | 1.20 |
| **DL-Methionine<sup>b</sup>** | 0.20 | 0.20 | 0.20 | 0.20 |
| **Chromium oxide<sup>g</sup>** | 1.00 | 1.00 | 1.00 | 1.00 |
| **Total** | 100.00 | 100.00 | 100.00 | 100.00 |

Fishmeal from wild catch (crude protein at 68%) was provided by IRIDA S.A, Greece.
<sup>b</sup> Provided by IRIDA S.A, Greece.
<sup>c</sup> Provided by Cooperative Farmers of Thessaly (THES-GI), Greece.
<sup>d</sup> Fish oil from wild catch was provided by IRIDA S.A, Greece.
<sup>e</sup> Per kilogram of mineral premix: 28 g Fe, 14 g Mn, 2.4 g I, 2.8 g Cu, 24 g Zn.
Provided by IRIDA S.A, Greece.
<sup>f</sup> Per kilogram of vitamin premix: 1200 mg retinol; 20 mg cholecalciferol; 400 mg biotin;
1.6 g folic acid; 60 g niacin; 24 g pantothenic acid; 8 g pyridoxine; 8 g riboflavin;
8 g thiamine; 80 mg vitamin B12; 80 g ascorbic acid; 100 g tocopherol acetate; 4 g

**Table 2.**
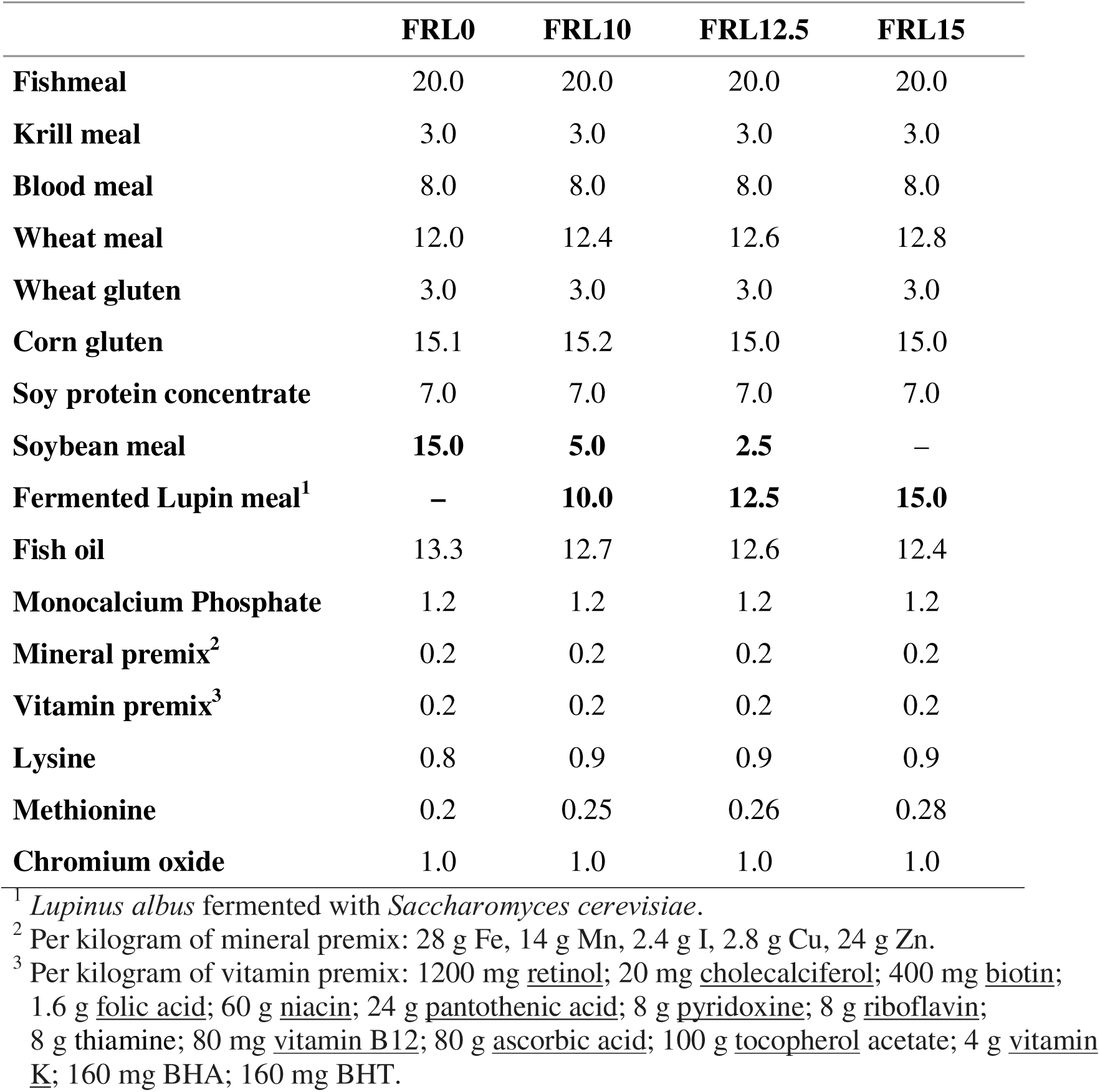
Formulation (% of wet weight) and composition of the Fermented *L. albus* feeding trial.

Detailed descriptions of the experimental design and methodology are provided in the studies by Vasilaki et al.(Vasilaki et al., 2025a, 2025b). In both feeding trials, a control diet containing soybean meal at an inclusion level of 15% was included for comparison. The experiments were conducted in a recirculating aquaculture system (RAS) using European sea bass (*D*. *labrax*) juveniles which were randomly allocated to 12 tanks (50 fish per tank) with each dietary treatment tested in triplicate. Fish fed *ad libitum* three times per day throughout the experimental period. The fermentation trial lasted 71 days and was initiated with fish of an average body weight of 18.9 ± 2.2 g, whereas the exogenous enzyme trial lasted 83 days and used fish with an initial average body weight at 11.2□ ±□ 1.3□ g.

The third feeding trial evaluated the use of *L. sceleratus* meal as fishmeal alternative in diets for *D*. *labrax.* Five experimental diets (Table 3) were formulated: a) a positive control diet containing 30% fishmeal (Control), b) a diet in which fishmeal was fully replaced by unprocessed *L. sceleratus* meal (UnP), included to assess the tolerance of European sea bass to tetrodotoxin (TTX), c) a diet in which *L. sceleratus* meal treated at high temperature replaced totally fishmeal (210°C / 10 min) and d) a diet in which *L. sceleratus* meal treated at low temperature replaced fully fishmeal (160°C /40 min) (T1H and T2L respectively), e) a diet in which 50% of fishmeal was replaced by T2L (Combo). The feeding trial lasted 106 days and was conducted in an open-flow seawater system. European sea bass juveniles with an initial average body weight 38.7 ± 4.3 g were randomly distributed in 15 tanks (25 fish/tank) providing three replicate tanks per dietary treatment. Fish were fed to apparent satiation twice daily throughout the experimental period.

**Table 3.** Formulation (% of wet weight) and composition of the *L. sceleratus* meal feeding trial. EPA: Eicosapentaenoic acid; DHA: Docosahexaenoic Acid; TTX: tetrodotoxin.

|  | **Control** |  | **UnP** |  | **T1H** | **T2L** | **Combo** |
| --- | --- | --- | --- | --- | --- | --- | --- |
| **Fishmeal** | 30 |  | - |  | - | - | 15 |
| **Lagomeal unprocessed** | - |  | 30 |  | - | - | - |
| **Lagomeal- High temperature** | - | - | 30 | - | - |  |  |
| **Lagomeal-Low temperature** | - |  | - |  | - | 30 | 15 |
| **Wheat meal** | **10.9** |  | **10.9** |  | **10.9** | **10.9** | **10.9** |
| **Corn gluten** | **19** |  | **19** |  | **19** | **19** | **19** |
| **Soybean Concentrate** | 17 | 17 | 17 | 17 | 17 |  |  |
| **Soybean meal** | 9 | 9 | 9 | 9 | 9 |  |  |
| **Fish oil** | 13 |  | 13 |  | 13 | 13 | 13 |
| **Monocalcium phosphate** | 0.5 | 0.5 | 0.5 | 0.5 | 0.5 |  |  |
| **Mineral and Vitamin premix** | 0.4 | 0.4 | 0.4 | 0.4 | 0.4 |  |  |
| **Lysine** | 0.1 | 0.1 | 0.1 | 0.1 | 0.1 |  |  |
| **Methionine** | 0.1 |  | 0.1 |  | 0.1 | 0.1 | 0.1 |
| **Proximate composition %** |  |  |  |  |  |  |  |
| **Protein** | 48 |  | 48 |  | 48 | 49 | 48 |
| **Fat** | 17 |  | 17 |  | 17 | 17 | 17 |
| **Moisture** | 6 |  | 8 |  | 6 | 7 | 6 |
| **Ash** | 7 |  | 8 |  | 8 | 8 | 8 |
| **EPA+DHA** | 2.3 |  | 1.9 |  | 1.8 | 1.7 | 2.1 |
| **TTX mg/kg** | 0 |  | 0.38 |  | 0 | 0 | 0 |

Diets were formulated and produced at the Applied Fish Nutrition Laboratory of the Institute of Marine Biology, Biotechnology and Aquaculture (Athens Greece). All experimental diets were designed to be isonitrogenous, isolipidic and isoenergetic, while meeting the nutritional requirements of European sea bass through balanced levels of essential amino acids, phosphorus, vitamins and minerals. Diets were produced using a twin-screw extruder (EV025A107FAA, Clextral, France), and oil was subsequently applied by vacuum coating using a Dinissen vacuum coater (The Netherlands).

### Sampling and DNA extraction

For the gut microbiota analysis, at the end of the feeding trials, fish were fasted for 24 hours. Then, nine fish per diet (3 fish/tank) were randomly collected, euthanized a lethal overdose of anaesthetic (a 1:10 clove oil and ethanol solution and their midguts were dissected under aseptic conditions, rinsed with sterile particle-free seawater and stored at −80°C until further analysis. Each midgut sample (n=117) was individually processed for DNA extraction using the QIAamp DNA Mini kit, following the manufacturer’s protocol for “DNA Purification from Tissues”.

### Amplicon sequencing and data analysis

Gut bacterial microbiota was characterised by 16S rRNA gene amplicon sequencing on Illumina MiSeq (2[×[300 bp) platform (Illumina, USA), targeting the V3-V4 region with the primer pair S-D-Bact-0341-b-S-17 and S-D-Bact-115 0785-a-A-21 (Klindworth et al., 2013). Data quality control and filtering performed in MOTHUR (v.1.48.0) (Schloss et al., 2009). The operational taxonomic units (OTUs) at 97% cutoff similarity level were taxonomically assigned with the Silva seed_v138_1 (Quast et al., 2013; Yilmaz et al., 2014). The presumptive metabolic functions of the found microbiota according to the 16S rRNA OTUs, were assessed by the PICRUSt2 (Douglas et al., 2020) software. All statistical analyses and plots were generated in RStudio (RStudio Team, 2015) using functions from the R packages phyloseq (McMurdie and Holmes, 2013), vegan (Oksanen et al., 2015), tidyverse (Wickham et al., 2019) and ggplot2 (Wickham, 2009).The raw data of this study have been submitted to the Short Read Archive (https://www.ncbi.nlm.nih.gov/sra) under the PRJNA1088722 accession number.

## RESULTS

A total of 4,014,404 sequence reads were clustered into 1,585 OTUs with 97% sequence identity. To ensure consistent sequencing depth across samples, reads were normalised at 5,645 reads/samples. As a result, nine (9) samples were eliminated from further analysis due to the low number of reads (Table 4), leaving the number of replicates per treatment between seven and nine. PERMANOVA analysis on the final sequencing data, with pairwise comparisons between individuals reared in replicate tanks within each dietary treatment, indicated that the gut microbiota did not differ among replicate tanks (P>0.05). These results indicate the absence of a tank effect and support the use of replicate tanks as biological replicates for subsequent comparison among dietary treatments

**Table 4.**
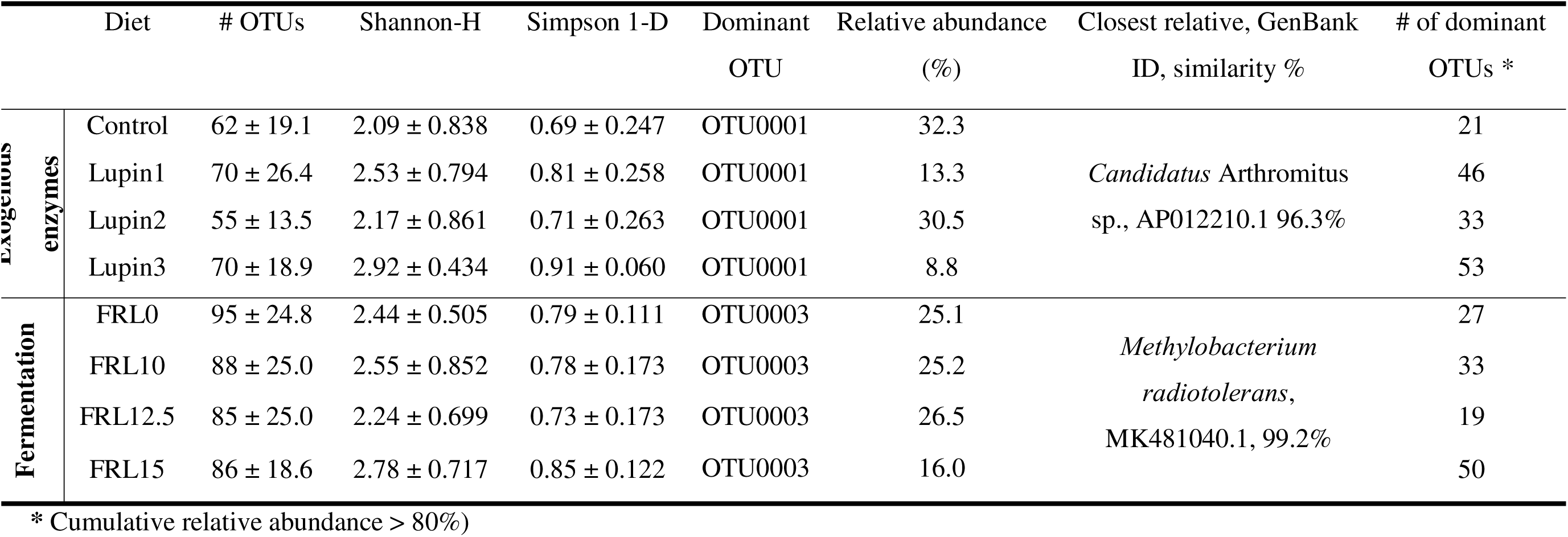
Alpha diversity results of the *D*. *labrax* midgut bacterial communities fed *L. albus* meal diets.

### *L. albus* meal trials

The gut microbiota of the European sea bass juveniles fed with *Lupinus albus* meal diets demonstrated high diversity, as reflected in both Simpson 1-D and Shannon-H indices (Table 4). The fermented *L*. *albus* diet had on average, 24 (37.7%) more observed taxa compared to the exogenous enzyme-treated feed. However, higher inclusion levels of *L*. *albus* meal in both trials led to an increased number (2.5 and 1.9 times more in exogenous enzyme and fermentation, respectively) of dominant OTUs compared to control (Table 4). PERMANOVA analysis based on the Bray-Curtis distance matrix, revealed that while the higher inclusion level of lupin meal led to greater variations in gut microbiome composition, these differences were not statistically significant when compared to the other inclusion levels of Lupin meal and the control diet. Furthermore, within each dietary group, including both fermentation and exogenous enzymes, there were no significant differences in gut microbiome composition between the different inclusion levels of *Lupinus* meal and the control diet. In contrast, significant differences (p < 0.05) in gut microbial compositions were found only between the two dietary groups (i.e. fermentation and exogenous enzymes) (Figure 1).

**Figure 1.**
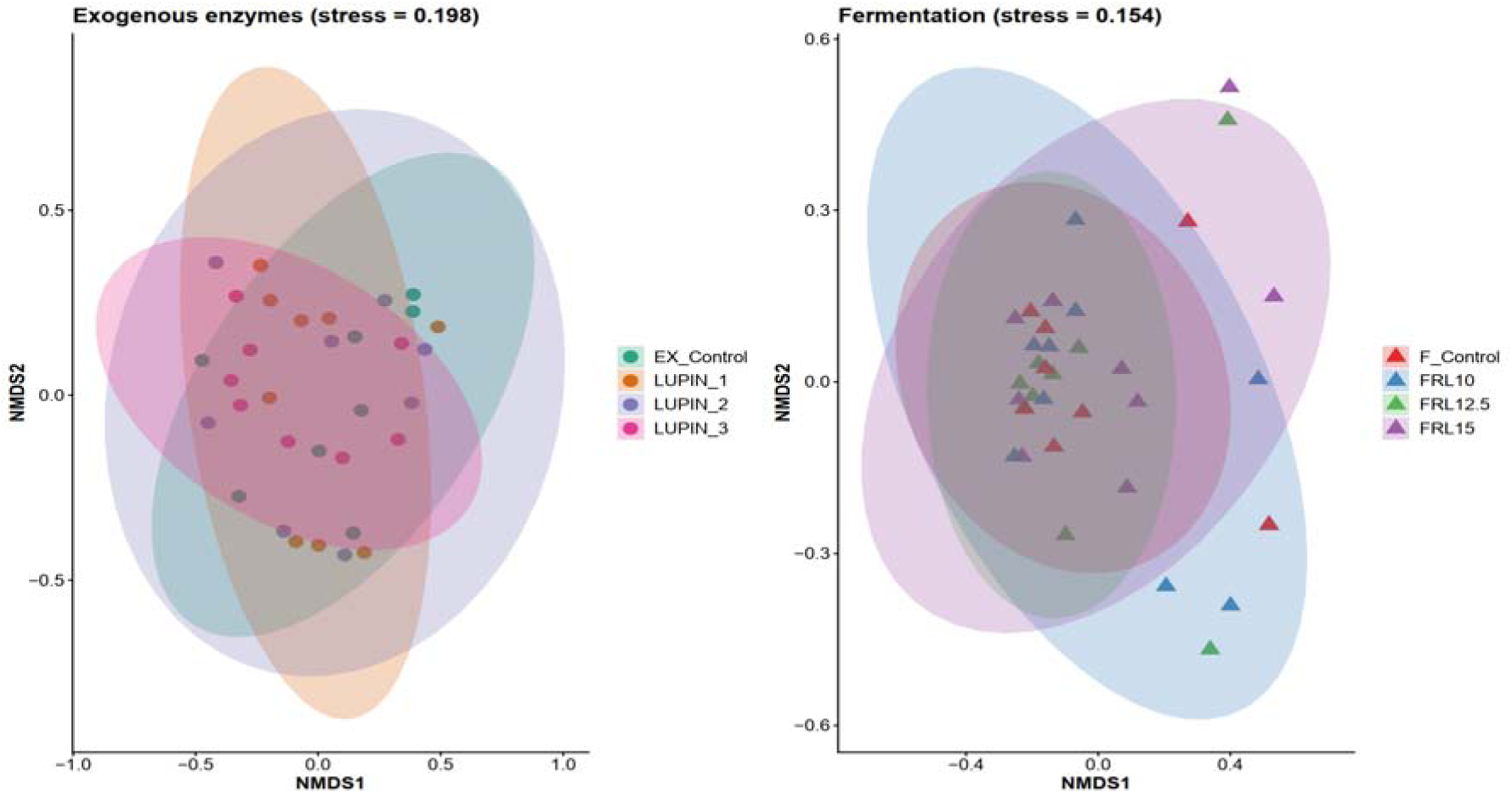
Non-metric multi-dimensional scaling of the lupin meal-fed *Dicentrarchus labrax* gut bacterial microbiota. FRL: fermented lupin meal.

Bacillota was identified as the dominant phylum in the gut microbial communities of the exogenous enzyme trial (Figure 2), with representatives of *Candidatus* Arthromitus (Clostridia) predominantly occurring across all diets (Figure 3, Table 4). In contrast, the fermentation trial displayed considerable dominance of Pseudomonadota as the dominant phylum (average Pseudomonadota/Bacillota ratio 5.9), accompanied by the prevalence of *Methylobacterium* at the genus level (Figure 2, Table 4).

**Figure 2.**
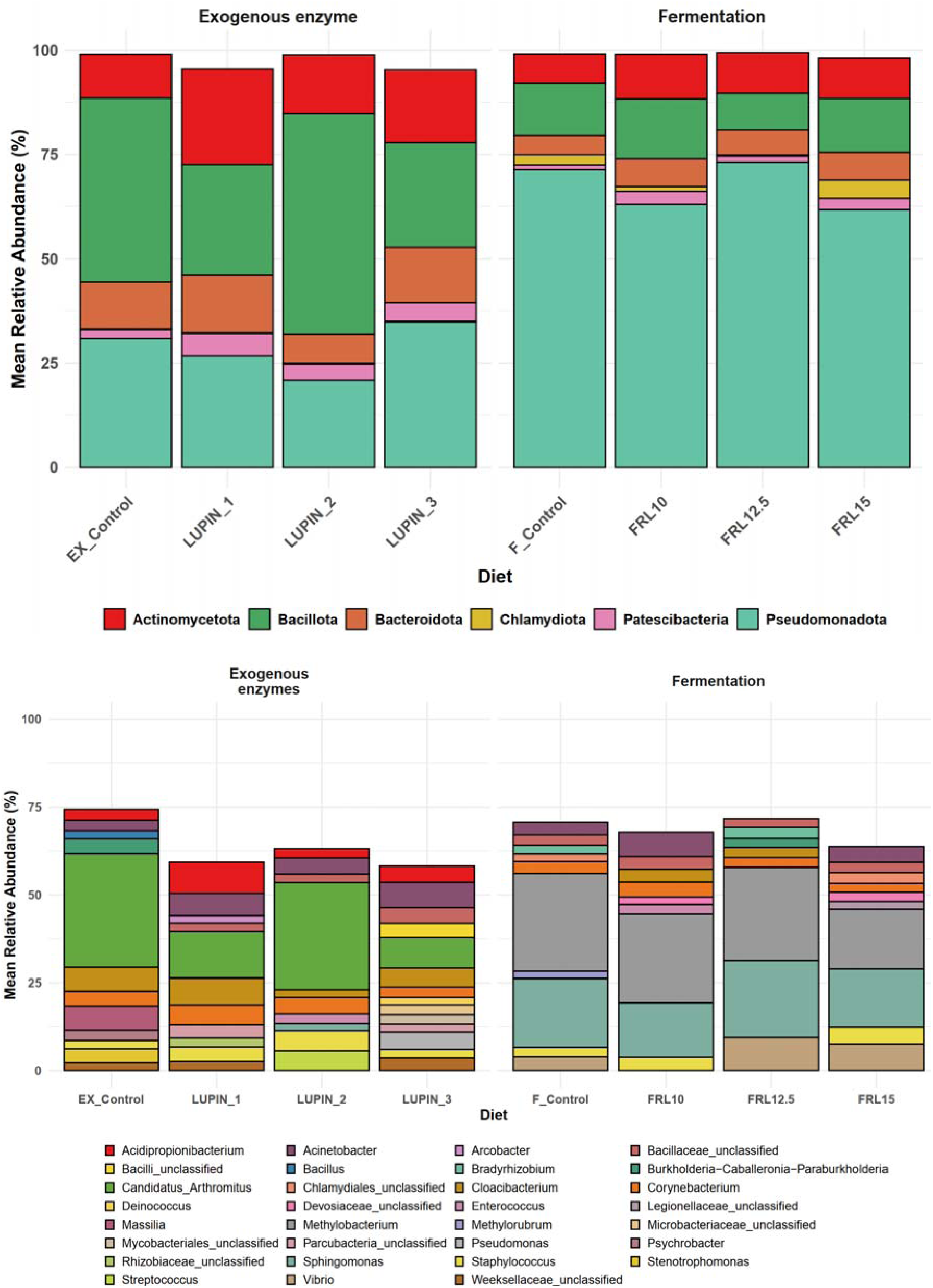
Most abundant phyla (top) and genera (bottom) gut microbiota of lupin meal-fed *Dicentrarchus labrax*. FRL: fermented lupin meal.

**Figure 3.**
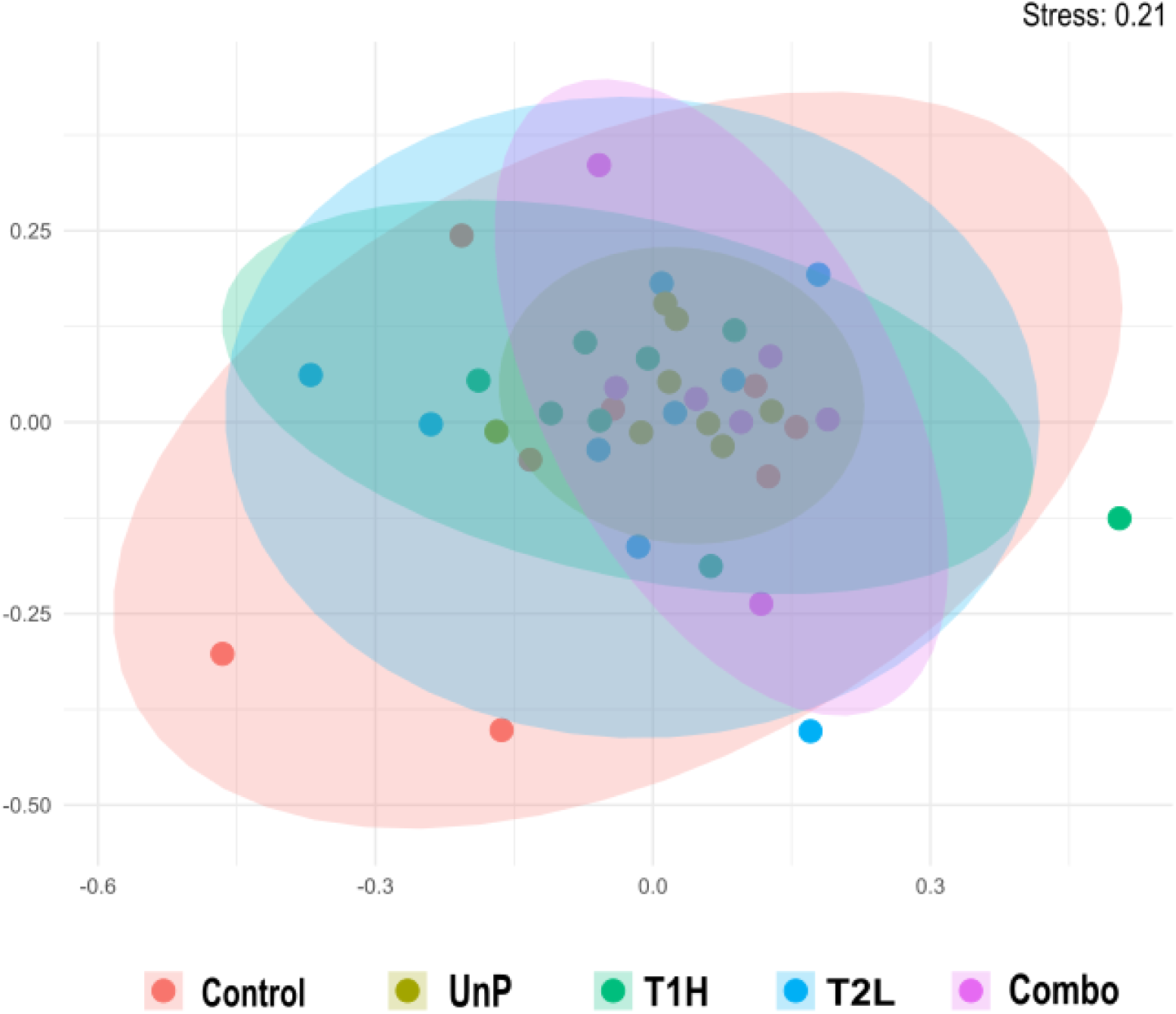
Non-metric multidimensional scaling of the *Lagocephalus sceleratus* meal-fed *Dicentrarchus labrax* gut microbiota.

### *L. sceleratus* meal

Feeding *D*. *labrax* with *L*. *sceleratus* meal diets, resulted in highly diverse gut bacterial microbiota, as reflected in both Simpson 1-D and Shannon-H diversity indices (Table 5). The group fed with the untreated-unprocessed *L*. *sceleratus* meal displayed the higher alpha diversity indices (Table 5). Although the observed variability in microbial composition, no significant differences were detected in the microbial distribution patterns between the groups of this feeding trial (PEMANOVA, p > 0.05, based on Bray-Curtis) (Figure 5).

**Table 5.** Alpha diversity results of the *D*. *labrax* midgut bacterial communities fed *L. sceleratus* meal.

|  | Taxa_S | Shannon-H | Simpson<br>1-D | Dominant OTU<br>(Relative abundance %) | Closest relative, GenBank<br>ID, similarity % | # of dominant<br>OTUs |
| --- | --- | --- | --- | --- | --- | --- |
| **Control** | 77 ± 23.3 | 3.03 ± 0.247 | 0.92 ± 0.03 | OTU0007<br>(6.7%) | *Staphylococcus hominis*,<br>MK212923, 99.8% | 59 |
| **UnP** | 102 ± 25.0 | 3.44 ± 0.202 | 0.94 ± 0.012 | OTU0007<br>(8.4%) | *Staphylococcus hominis*,<br>MK212923, 99.7% | 63 |
| **T1H** | 74 ± 11.6 | 2.86 ± 0.419 | 0.89 ± 0.061 | OTU0006<br>(14.0%) | *Cutibacterium acnes*,<br>CP025935, 98.8% | 47 |
| **T2L** | 73 ± 11.9 | 2.85 ± 0.349 | 0.89 ± 0.050 | OTU0006<br>(9.0%) | *Cutibacterium acnes*,<br>CP025935, 98.8% | 51 |
| **Combo (T2L**<br>**+Control)** | 83 ± 13.8 | 3.23 ± 0.172 | 0.93 ± 0.015 | OTU0009<br>(7.5%) | *Corynebacterium*<br>*vitaeruminis*, MW553688,<br>99.5% | 59 |

SIMPER analysis revealed differences in microbial community composition between the Control and the experimental treatments. The highest average dissimilarity was observed between the Control and the T2L (Low temperature) group (83.32%). In contrast, the highest similarity in microbial community composition was observed between the Control and the UnP control group (21.58%).

PERMANOVA analysis based on Bray–Curtis dissimilarities indicated that diet explained 16% of the total variation in gut microbial composition (R² = 0.162), but differences among dietary groups were not statistically significant (p = 0.453). NMDS ordination based on Bray–Curtis distances showed that samples from different dietary treatments did not form distinct clusters (Figure 3), indicating overlap in microbial community composition among diets.

The gut microbial communities in all diets of this trial were dominated by Pseudomonadota (32.9%), Actinomycetota (23.0%), Bacillota (20.9%) and Bacteroidota (15.3%) (Figure 4). The most abundant genera observed were *Burkholderia-Caballeronia-Paraburkholderia* (10.0), *Neobacillus* (8.64), *Acidipropionibacterium* (8.48) and *Cloacibacterium* (8.29) which has not been detected in Lago_HT treatment (Figure 6). Overall, both the dominant phyla and the remaining abundant genera were consistently observed across treatments, exhibiting only minor variations in relative abundance (Figure 6).

**Figure 4.**
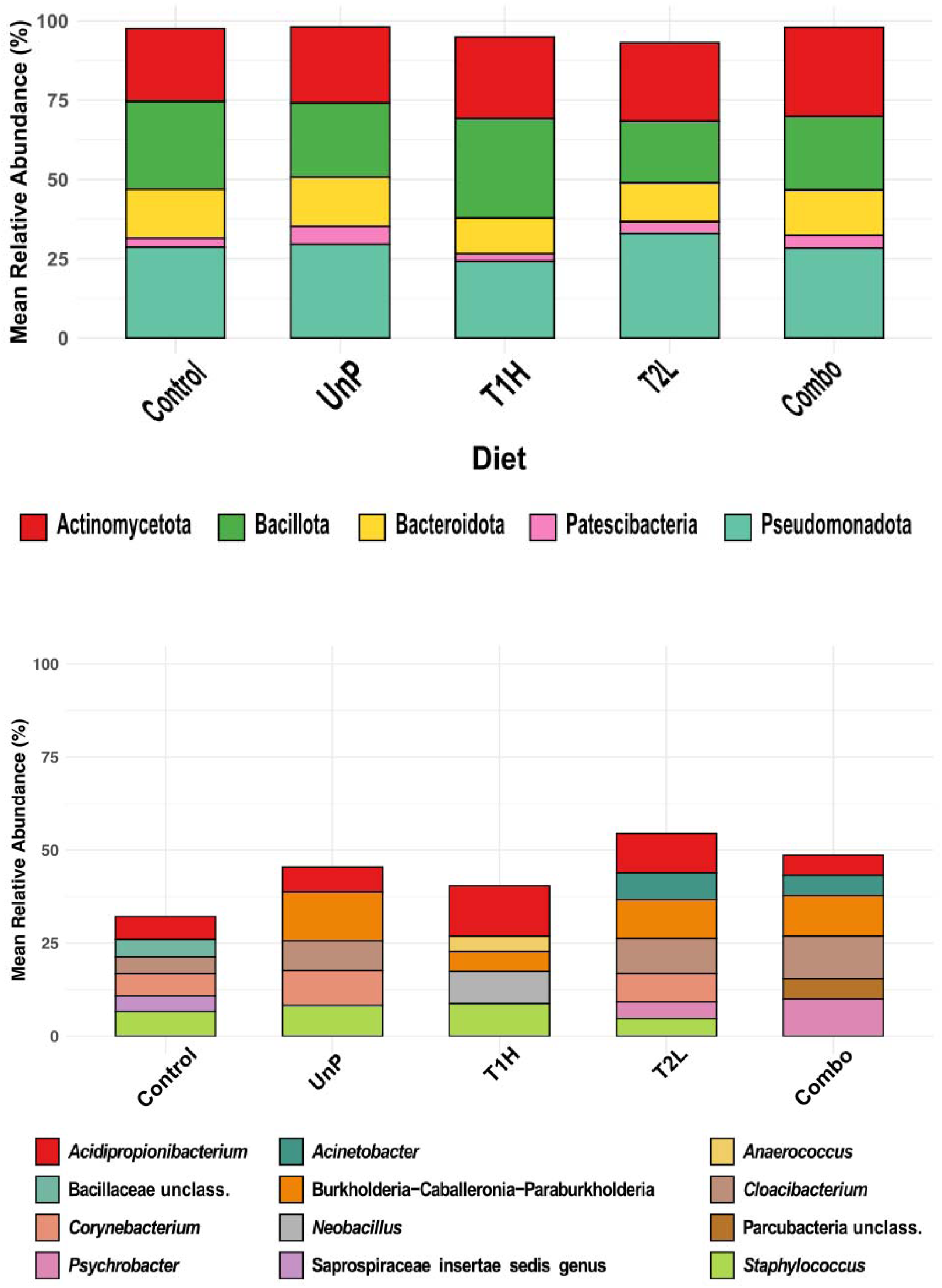
Most abundant phyla (top) and genera (bottom) gut microbiota of *Lagocephalus sceleratus* meal-fed *Dicentrarchus labrax*.

The Pseudomonadota/Bacillota ratio differed among treatments, ranging from 0.68 (T1H) to 6.70 (Compo). Control (1.03) and UnP diets (1.27) presented relatively balanced proportions, whereas T2L (2.95) showed a higher relative abundance of Bacillota compared to Pseudomonadota.

**Figure 5.**
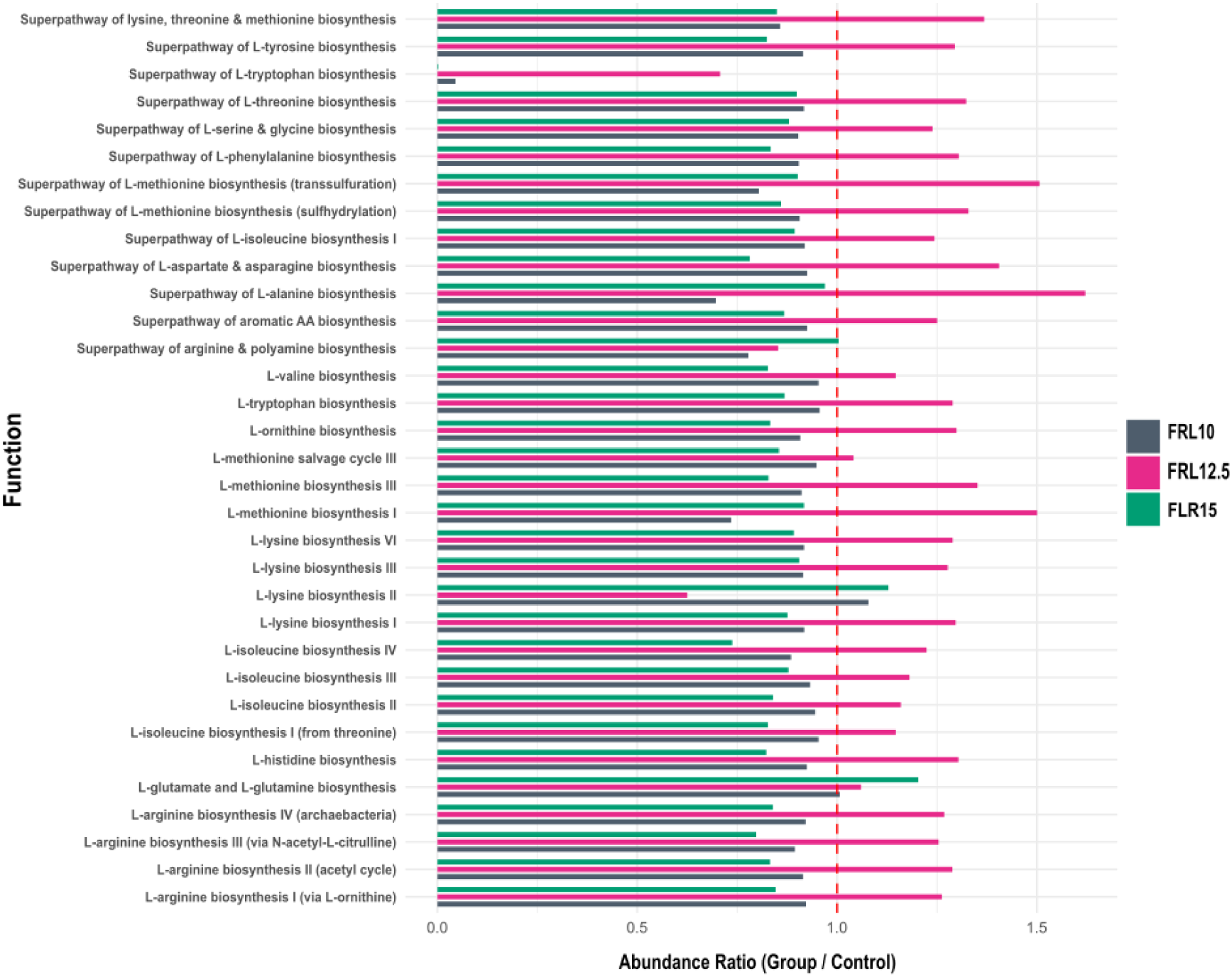
Major inferred gut bacterial metabolic pathways of lupin meal-fed *Dicentrarchus labrax*.

## DISCUSSION

The present study evaluated the effects of varying inclusion levels of novel sustainable dietary protein sources on the gut microbiota of *Dicentrarchus labrax* juveniles as potential aquafeed ingredients. Ideally a novel aquafeed is expected to either leave the gut microbiota rather unchanged compared to that of the standard aquafeed or enrich the fish gut habitat with beneficial bacteria. Two main ingredients were tested: *Lupinus albus* meal, processed through hydrolysis with exogenous enzymes or fermented with *Saccharomyces cerevisiae* to enhance nutrient bioavailability and digestibility, and heat-treated *Lagocephalus sceleratus* meal, to reduce tetrodotoxin (TTX) content. These were compared to conventional type diets based on soybean meal and commercial fishmeal. While direct studies on *D*. *labrax* gut microbiota fed diets with lupin are limited (Gatesoupe et al. 2014 & 2016), to our knowledge, the present study represents the first attempt to incorporate *L*. *sceleratus* meal into aquafeeds, marking a novel contribution to the exploration of alternative marine-based protein sources and also contribute to its invasive expansion.

### *L. albus* meal

No statistically significant changes were detected across different dietary inclusion levels or between the experimental diets (fermentation and hydrolysis with exogenous enzymes) and the control group, suggesting a moderate alteration of the gut microbiota of the *D. labrax* juveniles fed with the novel aquafeeds. The absence of significant gut bacterial community divergence of the treated animals from the control diet may be attributed to the processing of lupin meal, which likely reduced or eliminated the impact of common anti-nutritional factors, including non-starch polysaccharides (Abraham et al., 2019), alkaloids (Musco et al., 2017), and raffinose (Wolko et al. 2010). These compounds, if not mitigated, are known to interfere with nutrient absorption, damage intestinal epithelium, and alter microbial balance (Francis et al., 2001; Krogdahl et al., 2003). The hydrolysis and fermentation treatments applied here likely improved nutrient bioavailability and prevented potential dysbiotic effects, even at the higher inclusion level. However, in contrast to our findings, Gatesoupe et al. (2014), the only published study to date investigating the impact of lupin meal on the gut microbiota of *D*. *labrax*, reported significant alterations (p < 0.0001) in microbial composition following lupin inclusion. Their study assessed both mucosal and fecal microbial communities, with fecal samples showing the higher dissimilarity from the control group. These differences may be partially explained by the methodological different microbiota profiling approaches employed. Additionally, these may be attributed to the use of raw lupin meal (∼30%) in their diets, which is known to contain higher levels of anti-nutritional factors that can disrupt gut physiology and alter microbial communities. Furthermore, the use of fecal samples (i.e. transient Bacteria) and not gut tissue (i.e. more resident Bacteria) may have contributed to the observed differences between studies.

Moreover, in the present study, all key performance and somatometric indices did not differ significantly among treatments, with the only exception being the Lupin 1 group, which achieved a significantly higher final body weight (*P* = 0.050) compared to the control group (Vasilaki et al., 2025a, 2025b). All these results are aligned with previous research indicating that moderate inclusion levels of processed or raw lupin meal are generally well-tolerated by various fish species (Anwar et al., 2020; Hernández and Roman, 2016; Hughes, 1991; Salini and Adams, 2014). In the gilthead sea bream (*Sparus aurata*), the inclusion of up to 30% raw lupin meal has been shown to maintain growth performance at levels comparable to those obtained to fishmeal-based control diets (Pereira and Oliva-Teles, 2004). Furthermore, in the same study, micronised lupin meal significantly promoted higher growth rates, likely due to improved nutrient digestibility and reduced anti-nutritional factors, compared to raw ingredient.

Additionally, the gut microbiota of groups fed with higher inclusion levels of the hydrolysed and fermented lupin diets, became more heterogenous, resulting in an increased number of dominant OTUs (2.5x and 1.9x respectively, compared to the control group). This suggests that processing may encourage the proliferation of specific microbial taxa, but not enough to statistically separate inclusion levels into distinct clusters. Moreover, the fermented lupin diet supported a 37.7% more observed taxa than the enzyme-treated group, suggesting that fermentation not only alters nutrient bioavailability but also introduces microbial niches or metabolites that enrich microbial diversity. Similar effects have been observed in rainbow trout and carp when fed fermented plant proteins (Burel et al., 2000; Merrifield et al., 2010).

The high alpha diversity as reflected by both Shannon and Simpson indices, aligns with previous findings in teleosts fed high-fiber or legume-based diets, where plant-derived non-starch polysaccharides increase substrate complexity and promote microbial richness (Desai et al., 2012; Gajardo et al., 2017). High levels of lupin meal introduce large amounts of non-starch polysaccharides and other complex carbohydrates plus anti-nutritional factors into the diet. During enzymatic hydrolysis or fermentation complex macromolecules present in lupin meal are partially degraded into lower molecular weight compounds including oligo-and mono-saccharides, peptides and phenolic metabolites. The resulting increase in readily fermentable substrates and reduction of anti-nutritional factors creates new ecological niches and stimulatory compounds (Gullón et al., 2015), which together support a more varied and abundant set of dominant bacterial taxa.

Interestingly, *Ca*. Arthromitus sp. (OTU0001) and *Methylobacterium radiotolerans* (OTU0003) emerged as the dominant taxa in the exogenous enzyme and fermentation trials, respectively. These taxa were also prevalent in the corresponding control diets, suggesting that they represent core members of the gut microbiota in *D*. *labrax* under these dietary conditions. However, their relative abundances varied across treatments, indicating that diet processing may influence their proliferation or metabolic activity.

*C*α. Arthromitus [AP012210.1] (Prakash et al., 2011), is a non-cultivable member of the Segmented Filamentous Bacteria (SFB) and has been recently reclassified as *Anismomitus* genus (Kiran et al., 2026). SFB are well known for their close association with the intestinal epithelium, particularly in the ileum, where they play a critical role in modulating host immune responses by promoting the differentiation of Th17 cells. This immune activation has been linked to enhanced mucosal defences and reduced colonization by pathogenic bacteria (Gaboriau-Routhiau et al., 2009; Garland et al., 1982; Heczko et al., 2000). Despite their close relationship to pathogenic clostridia, SFB genomes lack major virulence factors, supporting their classification as beneficial immune modulators rather than pathogens. Their evolutionary divergence from an ancestral clostridial lineage, combined with adaptations for host-specific colonization, underscores their unique immunological role in promoting intestinal homeostasis (Klaasen et al., 1992; Schnupf et al., 2017, 2013). The detection of an OTU SFB-related in *D*. *labrax* here, along with a previous observation in the distal intestine of adult rainbow trout (*Oncorhynchus mykiss*) (Urdaci et al., 2001) and the finless porpoise (*Neophocaena asiaeorientalis asiaeorientalis*) feces and rectal swabs (Farkas et al. 2015, Ericsson et al. 2015), extending the known ecological and functional range of SFB beyond terrestrial vertebrates (Ericsson et al., 2015; Farkas et al., 2015), highlighting their potential significance (gut immune regulation and pathogen resistance) in aquatic host gut habitats as well.

In the fermented lupin meal trial, OTU0003 dominated and was affiliated with *Methylobacterium radiotolerans*. *Methylobacterium* species can utilize methanol, methane, and formic acid for growth and single cell protein production (Chistoserdova and Kalyuzhnaya, 2018; Patt et al., 1976). They also play a critical role in the carbon cycle by utilizing one-carbon compounds such as methanol and promoting plant growth (Wang et al., 2004; White et al., 2019). These bacteria are commonly found in a variety of natural environments, including plant leaf surfaces and soil (Sy et al., 2001), where they are known for their aerobic metabolism, often decomposing glucose oxidatively rather than fermentatively. Their presence have also been reported previously in high abundance within the autochthonous gut bacterial community in *D*. *labrax* (Carda-Diéguez et al., 2014; Pérez-Pascual et al., 2020; Rimoldi et al., 2020), and in the gut microbiome of other fish species (Kanika et al., 2025). 2025). Moreover, in brook charr (*Salvelinus fontinalis*), have shown a competitive relationship with *Flavobacterium* spp. (Boutin et al., 2014). Despite that may possess notable antimicrobial and cytotoxic properties (Balachandran et al., 2012), their role in the gut fish microbiome remains unclear and their presence in high abundances in sea bass gut microbiome merits further investigation. The beneficial features of *Methylobacterium* spp. are also depicted by their use as single-cell protein sources for animal feed, including fish feed (Gundupalli et al., 2024).

Despite the statistically insignificant differences between dietary trials, some noteworthy compositional changes took place. The fermentation group also showed a much higher Pseudomonadota/Bacillota ratio of 5.54 compared to the 0.79 of the enzyme-treated group, with dominance of *Methylobacterium-Methylorubrum*, while the enzyme-treated lupin group had a Bacillota-dominated microbiota with abundant Clostridia. These patterns suggest that fermentation may favour facultative anaerobes and bacteria capable of metabolizing fermentation byproducts (e.g., methanol, organic acids), whereas enzyme-treated diets may support obligate fermenters like unclassified Clostridia, known for their roles in short-chain fatty acid (SCFA) production.

### *L. sceleratus* meal

In this study, *L*. *sceleratus* meal inclusion was evaluated for the first time as a protein source in aquafeeds, and the results showed that, even though it’s a non-conventional protein source, it had limited effect on gut microbial structure. Moreover, the UnP group microbiota was more similar to the control (21.6%), suggesting that certain wild-caught or invasive species may be used without negative impacts at least in gut microbiome. However, the higher microbial diversity observed in unprocessed meal-fed fish (UnP group) may reflect a more complex substrate profile that promotes the proliferation of a broader range of microbial taxa (Gajardo et al., 2017; Huyben et al., 2019; Li et al., 2025).

Although the increased diversity could also be attributed to stress (Tolas et al., 2025), due to toxic components remaining in the unprocessed meal, according to Petersen and Round (2014) (Petersen and Round, 2014) dysbiosis is more commonly associated with a loss of diversity. Moreover, the dominant bacterial phyla observed in the gut microbial communities at *L*. *sceleratus* trial (Pseudomonadota, Bacillota, Actinomycetota and Bacteroidota), are commonly reported in the fish gut microbiomes (Spilsbury et al., 2022), and their consistent presence across all dietary treatments suggests a stable microbial structure. Moreover, at higher taxonomic resolution, 362 genera were detected across all dietary treatments, reflecting also a high degree of stability within the gut bacterial communities of *L*. *sceleratus*. Among these, *Acidipropionibacterium*, *Staphylococcus*, *Corynebacterium*, *Cloacibacterium*, *Burkholderia*-*Caballeronia*-*Paraburkholderia*, *Acinetobacter*, *Psychrobacter* as well as unclassified genera within the Bacillaceae and Parcubacteria, showed relative abundance >2%.

After performing BLAST analysis for these OTUs, some of the most abundant OTUs were found to have metabolic pathways important to their host. The OTUs related to *Methylobacterium fujisawaense* (OTU0003), *Sphingomonas phyllosphaerae* (OTU0004), *Cloacibacterium normanense* (OTU0005), *Cutibacterium acnes* (OTU0006), *Vibrio owensii* (OTU0008), *Corynebacterium vitaeruminis* (OTU0009) are able to synthesize lysine through the L-Aspartate pathway and arginine from citrulline. The enhancement of several amino acids biosynthesis was also shown especially for the FRL2 replacement by the metabolic inferred pathways for the whole communities (Figure 5). In addition, OTU0003 seems to have the full metabolic pathways for the biosynthesis of vitamins B1 (thiamine), B2 (riboflavin), B3 (nicotinate and nicotinamide), B5 (pantothenate), B7 (biotin) and B9 (folate). The observed enrichment in the specific metabolic pathways are more likely to represent an overexpression of the same microbiota as a result of increased metabolism, as has been reported for other animals’ symbionts e.g. (Cross et al., 2025), since no major changes were observed between the experimental treatments.

In conclusion, no significant differences were found in the gut bacterial microbiota of sea bass fed diets containing the pretreated sustainable ingredients, despite the differences in inclusion levels. This lack of significant impact on the microbiome could be attributed to the fact that the ingredients used in the diets were already pretreated through hydrolysis with exogenous enzymes, fermentation, or heat. These pretreatment methods seemed to have already optimized the ingredient for digestion and nutrient absorption, potentially reducing the need for further microbial modulation in the gut. As a result, variations in inclusion levels of the pretreated ingredients did not lead to a notable shift in the gut microbiome composition. However, moderate inclusion levels appeared to support a more balanced gut microbial community. Results from the *L*. *albus* suggest that fermented lupin with *Saccharomyces cerevisiae* may enhance nutrient availability and promote the growth of beneficial microbial populations. The apparent microbiota consistency presented in all feeding trials in this study, supports the potential of the tested ingredients as a novel protein source, contributing to sustainable feed diversification. Moreover, using *L*. *sceleratus* in aquafeeds simultaneously addresses sustainability (by reducing reliance on fishmeal and soybean meal) and invasive species control, aligning with circular economy principles. However, further validation under long-term and large-scale production conditions is required, mostly by functional metagenomic analyses to elucidate how these community changes translate into metabolic functions seem necessary to further elucidate the complete beneficial role of these sustainable aquafeed ingredients.

## ACKNOWLEDGEMENTS

This study is a part of two projects entitled “Improvement of the nutritional quality of forage legumes through biotechnological processes for the nutrition of Mediterranean aquaculture species” with MIS 5067490 and “An alternative way of utilizing the alien fish species of the genus Lagocephalus. Production of fishmeal for use in the diet of Mediterranean farmed marine species” with MIS 5067491, which have been co-funded by Greece and European Union, European Maritime and Fisheries Fund, in the context of the Operational Program “Maritime and Fisheries 2014–2020” (75[% EMFF contribution, 25[% National Contribution). This output reflects the views only of the author(s), and the European Union cannot be held responsible for any use which may be made of the information contained therein. The authors wish to thank Alltech for providing exogenous enzyme complex, Cooperative Farmers of Thessaly (THES-GI) for providing lupin seeds and IRIDA S.A for providing all other raw ingredients. The authors would also like to thank Eugenia Manolopoulou, Georgia Zoumpopoulou and Effie Tsakalidou from the Laboratory of Dairy Research, Agricultural University of Athens, Iera Odos 75 Athens, Greece for providing the *Saccharomyces cerevisiae* strain ACA-DC 5036 for the fermentation process.

## REFERENCES

Abraham, E.M., Ganopoulos, I., Madesis, P., Mavromatis, A., Mylona, P., Nianiou-Obeidat, I., Parissi, Z., Polidoros, A., Tani, E., Vlachostergios, D., 2019. The Use of Lupin as a Source of Protein in Animal Feeding: Genomic Tools and Breeding Approaches. Int. J. Mol. Sci. 20, 851. 10.3390/ijms20040851

Aimutis, W.R., Shirwaiker, R., 2024. A perspective on the environmental impact of plant-based protein concentrates and isolates. Proc. Natl. Acad. Sci. 121, e2319003121. 10.1073/pnas.2319003121

Anwar, A., Wan, A.H., Omar, S., El-Haroun, E., Davies, S.J., 2020. The potential of a solid-state fermentation supplement to augment white lupin (*Lupinus albus*) meal incorporation in diets for farmed common carp (*Cyprinus carpio*). Aquac. Rep. 17, 100348. 10.1016/j.aqrep.2020.100348

Balachandran, C., Duraipandiyan, V., Ignacimuthu, S., 2012. Cytotoxic (A549) and antimicrobial effects of Methylobacterium sp. isolate (ERI-135) from Nilgiris forest soil, India. Asian Pac. J. Trop. Biomed. 2, 712–716. 10.1016/S2221-1691(12)60215-9

Boutin, S., Sauvage, C., Bernatchez, L., Audet, C., Derome, N., 2014. Inter individual variations of the fish skin microbiota: host genetics basis of mutualism? PloS One 9, e102649. 10.1371/journal.pone.0102649

Burel, C., Boujard, T., Tulli, F., Kaushik, S.J., 2000. Digestibility of extruded peas, extruded lupin, and rapeseed meal in rainbow trout (*Oncorhynchus mykiss*) and turbot (*Psetta maxima*). Aquaculture 188, 285–298. 10.1016/S0044-8486(00)00337-9

Carda-Diéguez, M., Mira, A., Fouz, B., 2014. Pyrosequencing survey of intestinal microbiota diversity in cultured sea bass (*Dicentrarchus labrax*) fed functional diets. FEMS Microbiol. Ecol. 87, 451–459. 10.1111/1574-6941.12236

Chistoserdova, L., Kalyuzhnaya, M.G., 2018. Current Trends in Methylotrophy. Trends Microbiol. 26, 703–714. 10.1016/j.tim.2018.01.011

Coro, G., Vilas, L.G., Magliozzi, C., Ellenbroek, A., Scarponi, P., Pagano, P., 2018. Forecasting the ongoing invasion of *Lagocephalus sceleratus* in the Mediterranean Sea. Ecol. Model. 371, 37–49. 10.1016/j.ecolmodel.2018.01.007

Cross, K., Beckman, N., Jahnes, B., Sabree, Z.L., 2025. Microbiome metabolic capacity is buffered against phylotype losses by functional redundancy. Appl. Environ. Microbiol. 91, e02368–24. 10.1128/aem.02368-24

Darwis, Ridwanudin, A., Samsudin, R., Suryaningrum, L.H., 2026. Utilization of *Pterygoplichthys pardalis* as an alternative protein source in Nile tilapia feed: Effects on growth performance and health indicators. Egypt. J. Aquat. Res. 52, 128–136. 10.1016/j.ejar.2026.01.001

Darwis, Syahrul, Haerunnisa, Yusran, Sulfanita, A., Sufardin, Zulkarnaen, A., Yani, A., Yanti, N.D., 2025. Transforming an Invasive Species into Functional Feed: Growth, Nutritional Value, and Safety of Pterygoplichthys pardalis Meal for the Nile Tilapia. Egypt. J. Aquat. Biol. Fish. 29, 1925–1936. 10.21608/ejabf.2025.428024.6680

Desai, A.R., Links, M.G., Collins, S.A., Mansfield, G.S., Drew, M.D., Van Kessel, A.G., Hill, J.E., 2012. Effects of plant-based diets on the distal gut microbiome of rainbow trout (*Oncorhynchus mykiss*). Aquaculture 350–353, 134–142. 10.1016/j.aquaculture.2012.04.005

Douglas, G.M., Maffei, V.J., Zaneveld, J.R., Yurgel, S.N., Brown, J.R., Taylor, C.M., Huttenhower, C., Langille, M.G.I., 2020. PICRUSt2 for prediction of metagenome functions. Nat. Biotechnol. 38, 685–688. 10.1038/s41587-020-0548-6

Ericsson, A.C., Turner, G., Montoya, L., Wolfe, A., Meeker, S., Hsu, C., Maggio-Price, L., Franklin, C.L., 2015. Isolation of segmented filamentous bacteria from complex gut microbiota. BioTechniques 59, 94–98. 10.2144/000114319

Eroldoğan, O.T., Glencross, B., Novoveska, L., Gaudêncio, S.P., Rinkevich, B., Varese, G.C., de Fátima Carvalho, M., Tasdemir, D., Safarik, I., Nielsen, S.L., Rebours, C., Lada, L.B., Robbens, J., Strode, E., Haznedaroğlu, B.Z., Kotta, J., Evliyaoğlu, E., Oliveira, J., Girão, M., Vasquez, M.I., Čabarkapa, I., Rakita, S., Klun, K., Rotter, A., 2023. From the sea to aquafeed: A perspective overview. Rev. Aquac. 15, 1028–1057. 10.1111/raq.12740

Farkas, A.M., Panea, C., Goto, Y., Nakato, G., Galan-Diez, M., Narushima, S., Honda, K., Ivanov, I.I., 2015. Induction of Th17 cells by segmented filamentous bacteria in the murine intestine. J. Immunol. Methods 421, 104–111. 10.1016/j.jim.2015.03.020

Francis, G., Makkar, H.P.S., Becker, K., 2001. Antinutritional factors present in plant-derived alternate fish feed ingredients and their effects in fish. Aquaculture 199, 197–227. 10.1016/S0044-8486(01)00526-9

Gaboriau-Routhiau, V., Rakotobe, S., Lécuyer, E., Mulder, I., Lan, A., Bridonneau, C., Rochet, V., Pisi, A., De Paepe, M., Brandi, G., Eberl, G., Snel, J., Kelly, D., Cerf-Bensussan, N., 2009. The key role of segmented filamentous bacteria in the coordinated maturation of gut helper T cell responses. Immunity 31, 677–689. 10.1016/j.immuni.2009.08.020

Gajardo, K., Jaramillo-Torres, A., Kortner, T.M., Merrifield, D.L., Tinsley, J., Bakke, A.M., Krogdahl, Å., 2017. Alternative Protein Sources in the Diet Modulate Microbiota and Functionality in the Distal Intestine of Atlantic Salmon (Salmo salar). Appl. Environ. Microbiol. 83, e02615–16. 10.1128/AEM.02615-16

Garland, C.D., Lee, A., Dickson, M.R., 1982. Segmented filamentous bacteria in the rodent small intestine: Their colonization of growing animals and possible role in host resistance toSalmonella. Microb. Ecol. 8, 181–190. 10.1007/BF02010451

Gatesoupe, F.J., Huelvan, C., Le Bayon, N., Le Delliou, H., Madec, L., Mouchel, O., Quazuguel, P., Mazurais, D., Zambonino-Infante, J.L., 2016. The highly variable microbiota associated to intestinal mucosa correlates with growth and hypoxia resistance of sea bass, Dicentrarchus labrax, submitted to different nutritional histories. BMC Microbiol. 16, 1–13. 10.1186/s12866-016-0885-2

Gatesoupe, F.-J., Huelvan, C., Le Bayon, N., Sévère, A., Aasen, I.M., Degnes, K.F., Mazurais, D., Panserat, S., Zambonino-Infante, J.L., Kaushik, S.J., 2014. The effects of dietary carbohydrate sources and forms on metabolic response and intestinal microbiota in sea bass juveniles, *Dicentrarchus labrax*. Aquaculture 422–423, 47–53. 10.1016/j.aquaculture.2013.11.011

Gullón, P., Gullón, B., Tavaria, F., Vasconcelos, M., Gomes, A.M., 2015. In vitro fermentation of lupin seeds (Lupinus albus) and broad beans (Vicia faba): dynamic modulation of the intestinal microbiota and metabolomic output. Food Funct. 6, 3316–3322. 10.1039/c5fo00675a

Gundupalli, M.P., Ansari, S., Vital da Costa, J.P., Qiu, F., Anderson, J., Luckert, M., Bressler, D.C., 2024. Bacterial single cell protein (BSCP): A sustainable protein source from methylobacterium species. Trends Food Sci. Technol. 147, 104426. 10.1016/j.tifs.2024.104426

Gupta, S., Vera-Ponce de León, A., Kodama, M., Hoetzinger, M., Clausen, C.G., Pless, L., Verissimo, A.R.A., Stengel, B., Calabuig, V., Kvingedal, R., Skugor, S., Westereng, B., Harvey, T.N., Nordborg, A., Bertilsson, S., Limborg, M.T., Mørkøre, T., Sandve, S.R., Pope, P.B., Hvidsten, T.R., La Rosa, S.L., 2024. The need for high-resolution gut microbiome characterization to design efficient strategies for sustainable aquaculture production. Commun. Biol. 7, 1391. 10.1038/s42003-024-07087-4

Gupta, S.K., Fotedar, R., Foysal, M.J., Priyam, M., Siddik, M.A.B., Chaklader, M.R., Dao, T.T.T., Howieson, J., 2020. Impact of varied combinatorial mixture of non-fishmeal ingredients on growth, metabolism, immunity and gut microbiota of Lates calcarifer (Bloch, 1790) fry. Sci. Rep. 10, 17091. 10.1038/s41598-020-72726-9

Hachana, S., Abdallah, B.B., Rabeh, I., Harrath, A.H., Juarez-Facio, A.T., Cheraief, I., Joubert, O., Elcafsi, M., 2025. Fatty Acid Profiling and Nutritional Lipid Index Analysis of Portunus Segnis Meal as a Functional Fishmeal Substitute in Chelon Ramada and Oreochromis Niloticus. Chem. Afr. 8, 4437–4452. 10.1007/s42250-025-01439-1

Heczko, U., Abe, A., Finlay, B.B., 2000. Segmented filamentous bacteria prevent colonization of enteropathogenic Escherichia coli O103 in rabbits. J. Infect. Dis. 181, 1027–1033. 10.1086/315348

Hernández, A.J., Roman, D., 2016. Phosphorus and nitrogen utilization efficiency in rainbow trout (Oncorhynchus mykiss) fed diets with lupin (Lupinus albus) or soybean (Glycine max) meals as partial replacements to fish meal. Czech J. Anim. Sci. 61, 67–74. 10.17221/8729-CJAS

Hughes, S.G., 1991. Use of lupin flour as a replacement for full-fat soy in diets for rainbow trout (*Oncorhynchus mykis*). Aquaculture 93, 57–62. 10.1016/0044-8486(91)90204-K

Huyben, D., Vidaković, A., Werner Hallgren, S., Langeland, M., 2019. High-throughput sequencing of gut microbiota in rainbow trout (Oncorhynchus mykiss) fed larval and pre-pupae stages of black soldier fly (Hermetia illucens). Aquaculture 500, 485–491. 10.1016/j.aquaculture.2018.10.034

Islam, M.J., Essu, E.J., Hossain, Md.F., Islam, Md.A., Paul, S.K., Sumon, M.A.A., Bapary, M.A.J., Doan, H.V., 2026. Can Invasive Suckermouth Catfish, Hypostomus Plecostomus, Be Used as Fishmeal? A Potential Solution to Eradicate it from the Buriganga River, Bangladesh. Aquac. Fish Fish. 6, e70221. 10.1002/aff2.70221

Kanika, N.H., Liaqat, N., Chen, H., Ke, J., Lu, G., Wang, J., Wang, C., 2025. Fish gut microbiome and its application in aquaculture and biological conservation. Front. Microbiol. 15, 1521048. 10.3389/fmicb.2024.1521048

Kiran, S., Cruz, A.R., Daniau, A., Ma, B., Marbouty, M., Pipoli Da Fonseca, J., Legrand, A., Baudry, L., Cokelaer, T., Bensussan, M., Moya-Nilges, M., Garneau, J., Monot, M., Ochieng, J.B., Antonio, M., Tamboura, B., de Vos, W.M., Salonen, A., Hossain, J., Omore, R., Sow, S.O., Sansonetti, P.J., Ravel, J., Cerf-Bensussan, N., Schnupf, P., 2026. Segmented filamentous bacteria are worldwide human gut commensals. Nat. Commun. 10.1038/s41467-026-70010-4

Klaasen, H.L., Koopman, J.P., Poelma, F.G., Beynen, A.C., 1992. Intestinal, segmented, filamentous bacteria. FEMS Microbiol. Rev. 8, 165–180. 10.1111/j.1574-6968.1992.tb04986.x

Klindworth, A., Pruesse, E., Schweer, T., Peplies, J., Quast, C., Horn, M., Glöckner, F.O., 2013. Evaluation of general 16S ribosomal RNA gene PCR primers for classical and next-generation sequencing-based diversity studies. Nucleic Acids Res. 41, e1. 10.1093/nar/gks808

Krogdahl, Å., Bakke-McKellep, A. m., Baeverfjord, G., 2003. Effects of graded levels of standard soybean meal on intestinal structure, mucosal enzyme activities, and pancreatic response in Atlantic salmon (Salmo salar L.). Aquac. Nutr. 9, 361–371. 10.1046/j.1365-2095.2003.00264.x

Li, X., Wang, G., Fu, R., Zhu, X., Ren, P., Zhang, L., Ai, Q., Sun, Y., Wang, Z., 2025. Intestinal microbiota was closely related to feed efficiency of *Larimichthys crocea* fed two fishmeal-free diets. Aquaculture 594, 741367. 10.1016/j.aquaculture.2024.741367

Luna, G.M., Quero, G.M., Kokou, F., Kormas, K., 2022. Time to integrate biotechnological approaches into fish gut microbiome research. Curr. Opin. Biotechnol. 73, 121–127. 10.1016/j.copbio.2021.07.018

Malarvizhi, K., Kalaiselvan, P., Ranjan, A., 2024. Unlocking the potential of lupin as a sustainable aquafeed ingredient: a comprehensive review. Discov. Agric. 2, 43. 10.1007/s44279-024-00054-x

McMurdie, P.J., Holmes, S., 2013. phyloseq: An R Package for Reproducible Interactive Analysis and Graphics of Microbiome Census Data. PLOS ONE 8, e61217. 10.1371/journal.pone.0061217

Merrifield, D.L., Dimitroglou, A., Foey, A., Davies, S.J., Baker, R.T.M., Bøgwald, J., Castex, M., Ringø, E., 2010. The current status and future focus of probiotic and prebiotic applications for salmonids. Aquaculture 302, 1–18. 10.1016/j.aquaculture.2010.02.007

Musco, N., Cutrignelli, M.I., Calabrò, S., Tudisco, R., Infascelli, F., Grazioli, R., Lo Presti, V., Gresta, F., Chiofalo, B., 2017. Comparison of nutritional and antinutritional traits among different species (Lupinus albus L., Lupinus luteus L., Lupinus angustifolius L.) and varieties of lupin seeds. J. Anim. Physiol. Anim. Nutr. 101, 1227–1241. 10.1111/jpn.12643

Oficialdegui, F.J., Bedmar, S., Kouba, A., Vimercati, G., Roessink, I., Clavero, M., 2026. Demystifying invasivorism as a management strategy. Proc. Natl. Acad. Sci. 123, e2507779123. 10.1073/pnas.2507779123

Oksanen, J., Blanchet, F.G., Kindt, R., Legendre, P., Minchin, P., O’Hara, B., Simpson, G., Solymos, P., Stevens, H., Wagner, H., 2015. Vegan: Community Ecology Package. R Package Version 22-1 2, 1–2.

Panteli, N., Kousoulaki, K., Antonopoulou, E., Carter, C.G., Nengas, I., Henry, M., Karapanagiotidis, I.T., Mente, E., 2025. Which Novel Ingredient Should be Considered the “Holy Grail” for Sustainable Production of Finfish Aquafeeds? Rev. Aquac. 17, e12969. 10.1111/raq.12969

Patt, T.E., Cole, G.C., Hanson, R.S., 1976. Methylobacterium, a New Genus of Facultatively Methylotrophic Bacteria. Int. J. Syst. Evol. Microbiol. 26, 226–229. 10.1099/00207713-26-2-226

Pereira, T.G., Oliva-Teles, A., 2004. Evaluation of micronized lupin seed meal as an alternative protein source in diets for gilthead sea bream Sparus aurata L. juveniles. Aquac. Res. 35, 828–835. 10.1111/j.1365-2109.2004.01073.x

Pérez-Pascual, D., Estellé, J., Dutto, G., Rodde, C., Bernardet, J.-F., Marchand, Y., Duchaud, E., Przybyla, C., Ghigo, J.-M., 2020. Growth Performance and Adaptability of European Sea Bass (Dicentrarchus labrax) Gut Microbiota to Alternative Diets Free of Fish Products. Microorganisms 8, 1346.

Petersen, C., Round, J.L., 2014. Defining dysbiosis and its influence on host immunity and disease. Cell. Microbiol. 16, 1024–1033. 10.1111/cmi.12308

Piazzon, M.C., Ghosh, K., Ringø, E., Kokou, F., 2025. Chapter 17 - The importance of gut microbes for nutrition and health, in: Kumar, V. (Ed.), Feed and Feeding for Fish and Shellfish. Academic Press, pp. 575–637. 10.1016/B978-0-443-21556-8.00013-2

Prakash, T., Oshima, K., Morita, H., Fukuda, S., Imaoka, A., Kumar, N., Sharma, V.K., Kim, S.-W., Takahashi, M., Saitou, N., Taylor, T.D., Ohno, H., Umesaki, Y., Hattori, M., 2011. Complete genome sequences of rat and mouse segmented filamentous bacteria, a potent inducer of th17 cell differentiation. Cell Host Microbe 10, 273–284. 10.1016/j.chom.2011.08.007

Quast, C., Pruesse, E., Yilmaz, P., Gerken, J., Schweer, T., Yarza, P., Peplies, J., Glöckner, F.O., 2013. The SILVA ribosomal RNA gene database project: improved data processing and web-based tools. Nucleic Acids Res. 41, D590–D596. 10.1093/nar/gks1219

Ragaza, J.A., Hossain, M.S., Kumar, V., 2021. The Potential of Invasive Alien Fish Species as Novel Aquafeed Ingredients, in: Sustainable Aquafeeds. CRC Press.

Rimoldi, S., Gini, E., Koch, J.F.A., Iannini, F., Brambilla, F., Terova, G., 2020. Effects of hydrolyzed fish protein and autolyzed yeast as substitutes of fishmeal in the gilthead sea bream (Sparus aurata) diet, on fish intestinal microbiome. BMC Vet. Res. 16, 118. 10.1186/s12917-020-02335-1

RStudio Team, 2015. RStudio: Integrated Development for R. RStudio.

Salini, M.J., Adams, L.R., 2014. Growth performance, nutrient utilisation and digestibility by Atlantic salmon (*Salmo salar* L.) fed Tasmanian grown white (*Lupinus albus*) and narrow-leafed (*L. angustifolius*) lupins. Aquaculture 426–427, 296–303. 10.1016/j.aquaculture.2014.02.020

Schloss, P.D., Westcott, S.L., Ryabin, T., Hall, J.R., Hartmann, M., Hollister, E.B., Lesniewski, R.A., Oakley, B.B., Parks, D.H., Robinson, C.J., Sahl, J.W., Stres, B., Thallinger, G.G., Van Horn, D.J., Weber, C.F., 2009. Introducing mothur: open-source, platform-independent, community-supported software for describing and comparing microbial communities. Appl. Environ. Microbiol. 75, 7537–7541. 10.1128/AEM.01541-09

Schnupf, P., Gaboriau-Routhiau, V., Cerf-Bensussan, N., 2013. Host interactions with Segmented Filamentous Bacteria: An unusual trade-off that drives the post-natal maturation of the gut immune system. Semin. Immunol., Microbiota and the immune system, an amazing mutualism forged by co-evolution 25, 342–351. 10.1016/j.smim.2013.09.001

Schnupf, P., Gaboriau-Routhiau, V., Sansonetti, P.J., Cerf-Bensussan, N., 2017. Segmented filamentous bacteria, Th17 inducers and helpers in a hostile world. Curr. Opin. Microbiol., Host-microbe interactions: bacteria 35, 100–109. 10.1016/j.mib.2017.03.004

Serra, V., Pastorelli, G., Tedesco, D.E.A., Turin, L., Guerrini, A., 2024. Alternative protein sources in aquafeed: Current scenario and future perspectives. Vet. Anim. Sci. 25, 100381. 10.1016/j.vas.2024.100381

Silva, F.C. de P., Nicoli, J.R., Zambonino-Infante, J.L., Kaushik, S., Gatesoupe, F.-J., 2011. Influence of the diet on the microbial diversity of faecal and gastrointestinal contents in gilthead sea bream (Sparus aurata) and intestinal contents in goldfish (Carassius auratus). FEMS Microbiol. Ecol. 78, 285–296. 10.1111/j.1574-6941.2011.01155.x

Spilsbury, F., Foysal, M.J., Tay, A., Gagnon, M.M., 2022. Gut Microbiome as a Potential Biomarker in Fish: Dietary Exposure to Petroleum Hydrocarbons and Metals, Metabolic Functions and Cytokine Expression in Juvenile Lates calcarifer. Front. Microbiol. 13, 827371. 10.3389/fmicb.2022.827371

Sy, A., Giraud, E., Jourand, P., Garcia, N., Willems, A., de Lajudie, P., Prin, Y., Neyra, M., Gillis, M., Boivin-Masson, C., Dreyfus, B., 2001. Methylotrophic Methylobacterium bacteria nodulate and fix nitrogen in symbiosis with legumes. J. Bacteriol. 183, 214–220. 10.1128/JB.183.1.214-220.2001

Szczepański, A., Adamek-Urbańska, D., Kasprzak, R., Szudrowicz, H., Śliwiński, J., Kamaszewski, M., 2022. Lupin: A promising alternative protein source for aquaculture feeds? Aquac. Rep. 26, 101281. 10.1016/j.aqrep.2022.101281

Tolas, I., Zhou, Z., Zhang, Z., Teame, T., Olsen, R.E., Ringø, E., Rønnestad, I., 2025. A fishy gut feeling – current knowledge on gut microbiota in teleosts. Front. Mar. Sci. 11. 10.3389/fmars.2024.1495373

Urdaci, M.C., Regnault, B., Grimont, P.A., 2001. Identification by in situ hybridization of segmented filamentous bacteria in the intestine of diarrheic rainbow trout (Oncorhynchus mykiss). Res. Microbiol. 152, 67–73. 10.1016/s0923-2508(00)01169-4

Vagenas, G., Karachle, P.K., Oikonomou, A., Stoumboudi, M.T., Zenetos, A., 2024. Decoding the spread of non-indigenous fishes in the Mediterranean Sea. Sci. Rep. 14, 6669. 10.1038/s41598-024-57109-8

Vasilaki, A., Nengas, I., Fountoulaki, E., Henry, M., Kogiannou, D., Nikoloudaki, C., Chronopoulos, P., Karapanagiotidis, I.T., Mente, E., 2025a. Nutritional enhancement of lupin meal (Lupinus albus), through fermentation with Saccharomyces cerevisiae, as plant protein ingredient in aquafeeds for the European sea bass (Dicentrarchus labrax). Aquaculture 609, 742820. 10.1016/j.aquaculture.2025.742820

Vasilaki, A., Nengas, I., Kogiannou, D., Henry, M., Nikoloudaki, C., Chronopoulos, P., Berillis, P., Golomazou, E., Fountoulaki, E., Karapanagiotidis, I.T., Mente, E., 2025b. Evaluation of the nutritional value of processed lupin meal (Lupinus albus) with exogenous enzymes, as feed ingredient in European sea bass (Dicentrarchus labrax) aquafeeds. Aquac. Rep. 45, 103135. 10.1016/j.aqrep.2025.103135

Wang, Y., Stingl, U., Anton-Erxleben, F., Zimmer, M., Brune, A., 2004. ‘Candidatus Hepatincola porcellionum’ gen. nov., sp. nov., a new, stalk-forming lineage of Rickettsiales colonizing the midgut glands of a terrestrial isopod. Arch. Microbiol. 181, 299–304. 10.1007/s00203-004-0655-7

White, J.F., Kingsley, K.L., Zhang, Q., Verma, R., Obi, N., Dvinskikh, S., Elmore, M.T., Verma, S.K., Gond, S.K., Kowalski, K.P., 2019. Review: Endophytic microbes and their potential applications in crop management. Pest Manag. Sci. 75, 2558–2565. 10.1002/ps.5527

Wickham, H., 2009. ggplot2: Elegant Graphics for Data Analysis, Use R! Springer-Verlag, New York.

Wickham, H., Averick, M., Bryan, J., Chang, W., McGowan, L.D., François, R., Grolemund, G., Hayes, A., Henry, L., Hester, J., Kuhn, M., Pedersen, T.L., Miller, E., Bache, S.M., Müller, K., Ooms, J., Robinson, D., Seidel, D.P., Spinu, V., Takahashi, K., Vaughan, D., Wilke, C., Woo, K., Yutani, H., 2019. Welcome to the Tidyverse. J. Open Source Softw. 4, 1686. 10.21105/joss.01686

Yilmaz, P., Parfrey, L.W., Yarza, P., Gerken, J., Pruesse, E., Quast, C., Schweer, T., Peplies, J., Ludwig, W., Glöckner, F.O., 2014. The SILVA and “All-species Living Tree Project (LTP)” taxonomic frameworks. Nucleic Acids Res. 42, D643–648. 10.1093/nar/gkt1209

Zhang, X., Ying, C., Jiang, M., Lin, D., You, L., Yin, D., Zhang, J., Liu, K., Xu, P., 2022. The bacteria of Yangtze finless porpoise (Neophocaena asiaeorientalis asiaeorientalis) are site-specific and distinct from freshwater environment. Front. Microbiol. 13, 1006251. 10.3389/fmicb.2022.1006251

Zhang, Z., Yang, Q., Liu, H., Jin, J., Yang, Y., Zhu, X., Han, D., Zhou, Z., Xie, S., 2025. Potential Functions of the Gut Microbiome and Modulation Strategies for Improving Aquatic Animal Growth. Rev. Aquac. 17, e12959. 10.1111/raq.12959

